# Bacterial Sensitivity to Chlorhexidine and Povidone-Iodine Antiseptics Over Time: A Systematic Review and Meta-Analysis of Human-Derived Data

**DOI:** 10.1101/2021.11.23.469660

**Authors:** Raiyyan Aftab, Vikash H Dodhia, Christopher Jeanes, Ryckie G Wade

## Abstract

**Background:** Surgical site infection (SSI) is the most common complication of surgery, increasing healthcare costs and hospital stay. Topical biocides such as chlorhexidine (CHX) and povidone-iodine (PVI) are used for skin antisepsis to minimise SSIs. There is an increasing concern of developing resistance to topical biocides, however the clinical implications of this remains unclear.

**Outcomes:** The objective of this review was to determine whether the Minimum Bactericidal Concentration (MBC) for topical preparations of CHX or PVI have changed over time, in microbes relevant to SSI.

**Methods:** We searched for studies which reported the mean bactericidal concentration (MBC) of laboratory and clinical isolates of common SSI causing microbes to CHX and PVI. We excluded samples derived from non-humans and studies using antimicrobial solvents or mixtures of biocides with other active substances. MBC was pooled in random effects meta-analyses and change in MBC over time was explored using meta-regression.

**Results:** 79 studies were including, analysing 6218 microbes between 1976 and 2021. Most studies used CHX (93%) and there was insufficient data for meta-analysis of PVI. Enterobacteriales had the highest MBC for CHX (20 mg/L [95% CI 14, 25]; I^2^ 95%) whilst MRSA had the lowest (3 mg/L [95% CI 1, 2]; I^2^ 93%). There was no change in MBC of CHX to *Staphylococci* (β 0.12 [-1.13, 1.37]; I^2^ 99%) or *Streptococci* (β 0.13 [-0.35, 0.62]; I^2^ 97%).

**Conclusions:** There is no evidence of change in susceptibility of common SSI-causing microbes to CHX over time. This study provides reassurance that the worldwide guidance that CHX should remain the first-choice agent for skin asepsis prior to surgery.

## Introduction

Surgical site infection (SSI) is the most common and costly complication of surgery(1,2), occurring in approximately 5% of all surgical interventions(3). They represent an important economic burden across all surgical specialties(4), increasing hospital inpatient stay time and adversely affecting patient’s mental and physical health.(4) The use of skin antisepsis prior to surgery significantly reduces the risk of SSI and consequently, post-operative morbidity and mortality(5–7).

*Staphylococcus aureus* and *Streptococc*i spp. are commonly implicated microbes in SSI, along with *Enterococcus* spp. and *Escherichia coli*(8). To reduce the risk of SSI, the World Health Organisation (WHO)(7), United States of America Centre for Disease Control (CDC)(9) and United Kingdom National Institute for Health and Care Excellence (NICE)(10) recommend the application of topical chlorhexidine (CHX) in alcohol to the planned operative site, for skin antisepsis. CHX in an alcoholic solvent has been shown to halve the risk of SSI following clean(11), contaminated(12,13) and dirty surgery when compared to other antiseptics such as povidone-iodine (PVI).

CHX is a biguanide compound, utilised both as a broad spectrum antimicrobial and topical antiseptic(14). By binding to the cell membrane and cell wall of bacteria, at lower concentrations it has a bacteriostatic effect by displacing the cations and destabilising the cell well. At higher concentrations there is a complete loss of cellular structural integrity, having a bactericidal effect.(15) PVI is an iodophor; a chemical complex between a water soluble povidone polymer and iodine(16). When dissolved in water, iodine is released, penetrating microorganisms and oxidising proteins, nucleotides and fatty acids(17) causing cell death. Both PVI and CHX are active against gram positive and negative bacteria, fungi and viruses(16,18).

There are growing fears that as antiseptic use increases, sensitivity may reduce and resistance emerge(19). This is particularly concerning given the accelerating global antibiotic resistance crisis(20). Multiple bacteria have shown reduced sensitivity (perhaps even resistance) to CHX, particularly through multidrug resistance efflux protein qacA(19). Methicillin resistant *Staphylococcus aureus* (MRSA) samples with qacA/B genes showed persistent MRSA carriage despite de-colonisation therapy(21). Not only is the presence of reduced sensitivity concerning, but the presence of resistance conferring genes seems to be increasing annually(22). PVI resistance has been less common reported(23) although this might be secondary to the multimodal effect of iodine on microbes(23) or otherwise. Overall, the current state of microbial sensitivity to CHX and PVI remains unclear.

The aim of this review is to summarise the sensitivity profiles of skin microbes (relevant to surgical site infection) to CHX and PVI, and explore how these have changed over time

## Methods

This review was designed and conducted in accordance with the Cochrane Handbook of Systematic Reviews(24), the protocol was published in the PROSPERO databased (CRD42021241089) and the report has been authored in accordance with the PRISMA checklist(25).

### Types of Studies

We included all studies which reported the resistance of microbes to CHX or PVI based topical biocides derived from human samples. There were no language restrictions. We excluded case reports and studies which used antimicrobial solvents (e.g., alcohol) or mixtures of antiseptics (e.g. chlorhexidine mixed with cetrimide).

### Search strategy

The NICE Healthcare Databases (hdas.nice.org.uk) was searched according to appendix 1 (supplementary materials). The medRxiv and bioRxiv preprint archives were searched with the same strategy using the R package medrxivr(26). This yielded 582 hits in PubMed, 720 in Embase, 993 in Web of Science and 896 in CINAHL. After de-duplication, there were 2318 unique citations which were screened (Fig 1). A further 3 articles were found by manual searching of these articles.

**Fig 1.**
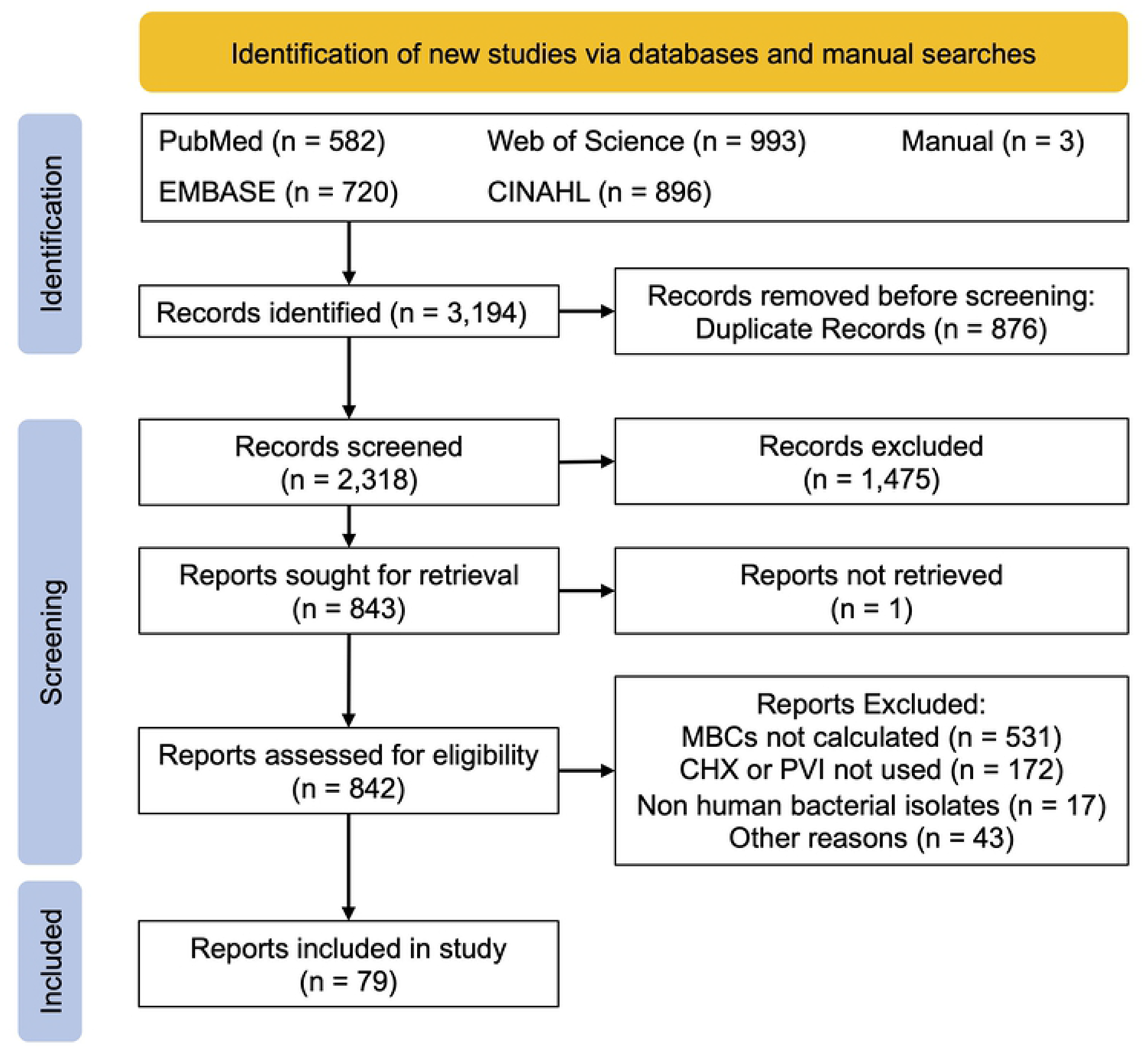
PRISMA flowchart

### Study selection

Two review authors (RA and VHD) independently screened titles and abstracts for relevance, in accordance with the eligibility criteria. The full texts of potentially eligible articles were obtained and again independently assessed by the same authors. Disagreements were resolved by discussion with RGW.

### Data extraction

Two review authors (RA and VHD) independently double extracted data. The colony was the unit of analysis. Where data was missing or unclear, the corresponding author was contacted by email and if no reply was received, these values were estimated from the available data(27).

### Outcomes

The outcome of interest was the Minimum Bactericidal Concentration (MBC). MBC was chosen over the minimum inhibitory concentration (MIC) as a measure of susceptibility because it is generally felt to be a more appropriate for topical antiseptics(15,28). We were interested in deriving a pooled estimate of the MBC for different species to understand how this may have changed over time.

### Methodological quality assessment

The risk of bias and methodological quality was not assessed because there are no validated tools available for studies of this nature and the study selection process ensured that only the highest quality research was included.

### Missing data

After estimating some parameters, the overall rate of missing data (in the predictor variables) was 14.5%. Importantly, the variance of the mean MBC was missing at random in 36 observations (61%) and because this was required for the primary analyses, we imputed this data using chained equations(29,30).

### Statistical analysis

The raw data are available open-source at https://osf.io/khnb2. Using metafor(31,32), 5 studies(33–37) reporting the MBC of CHX for *Staphylococci* were identified as outliers (based on externally standardised residuals) or influential studies (based on the Cook’s distance and leave-one-out values of the test statistics for heterogeneity), and their 95% confidence intervals (CIs) were far outside the 95% CI of the pooled estimate, so they were excluded from the meta-analyses. Data were then analysed in Stata/MP v16 (StataCop LLC, Texas) using the meta suite. To synthesise a pooled MBC, we used a random-effects meta-analyses. We sub-grouped by the microbial family. To estimate change in MBC over time, we performed meta-regression for *Staphylococci* and *Streptococci*, separately. A sensitivity multivariable meta-regression for *Staphylococci* was performed, controlling for methicillin-resistance as a binary co-variate. The REML estimator was used throughout. To align with calls for the abolition of p-values, we minimise their use and avoid the term “statistical significance”(38,39), instead focusing on how our findings may be clinically applicable and what might explain uncertainty in the estimates.

## Results

Ultimately, 79 studies(33–37,40–113) were included (Fig 1).

### Study characteristics

The details of the included studies are summarised in S1 Table; readers who wish to know more detail should refer to the raw data (https://osf.io/khnb2/). The included studies originated from 24 countries. Articles were published between 1976 and 2021, although the majority (95%) were published this century. The antiseptics used included five different CHX salts (digluconate(33,35,37,40,47,50,55–58,61,63,67,69,74,75,81,83,88,90,94,97,99,101,106,107,111,113), gluconate(34,36,42,42,53,91,102,103,108,108,109,112), dichlorohydrate(85–87), diacetate(71,99) and dihydrochloride(96)) and povidone-iodine(37,46,49,70,76,91). In total, MBC data from 6218 microbes were extracted. The microbes tested are shown in S2 Table. Most samples were laboratory isolates (61%) and not multi-drug resistant (88%). The reporting standards used to establish the MBC were the according to the Clinical Laboratory Standards Institute (CLSI, 67%), European Committee on Antimicrobial Testing (EUCAST, 6%), German Institute for Standardisation (DIN, 5%), British Society for Antimicrobial Chemotherapy (BSAC, 3%) or International Organisation for Standardisation (ISO, 1%).

### Evidence Synthesis

The MBC of CHX differed significantly between the families of microbes (Fig 2). Enterobacteriales had the highest MBC for CHX (20 mg/L [95% CI 14, 25]; I^2^ 95%) whilst MRSA had the lowest (3 mg/L [95% CI 1, 2]; I^2^ 93%).

**Fig 2.**
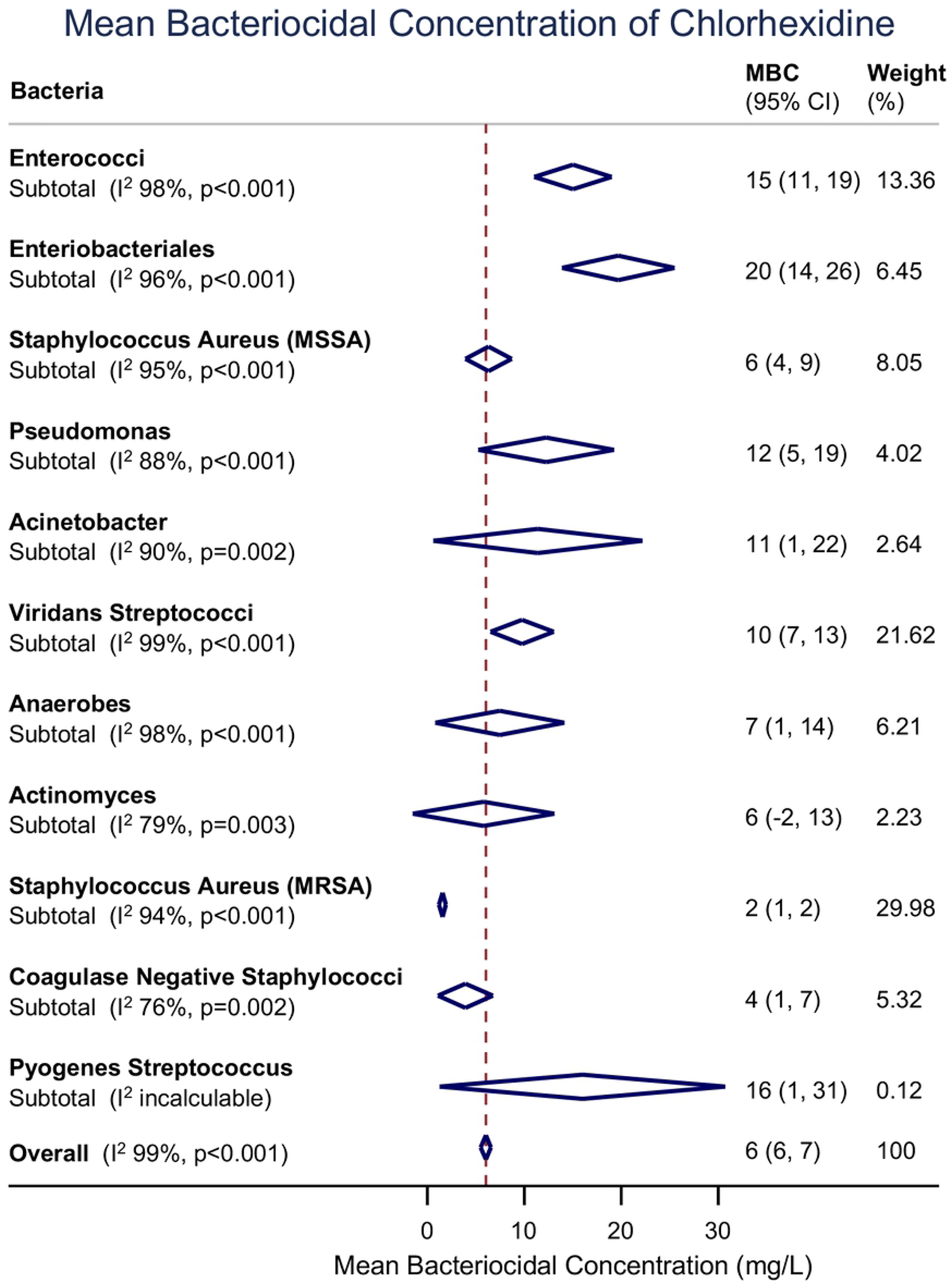
Forest plot of the mean MBC for different species and families of bacteria

Overall, 23 studies reported the mean MBC for *Staphylococci*; observations based on MSSA were more common(41,44,51,59,61,67,68,74,88,93–95,102,106,107,110–113) than MRSA(34,35,37,50,67,68,80,90,93,106,110,113). The pooled mean MBC of CHX for *Staphylococci* was 5.93 mg/L (95% CI 3.09, 8.77; I2 99%). Meta-regression showed no change in the MBC of CHX for *Staphylococci* over time (β 0.12 [-1.13, 1.37]; I^2^ 99%; Fig 3). When controlling for resistance to methicillin (MRSA vs MSSA), there was still no evidence of a change in the MBC over time (β 0.26 [-0.87, 1.34]; I^2^ 99%). Study level estimates for MSSA, MRSA and coagulase-negative *Staphylococci* are shown in S1 Fig.

**Fig 3.**
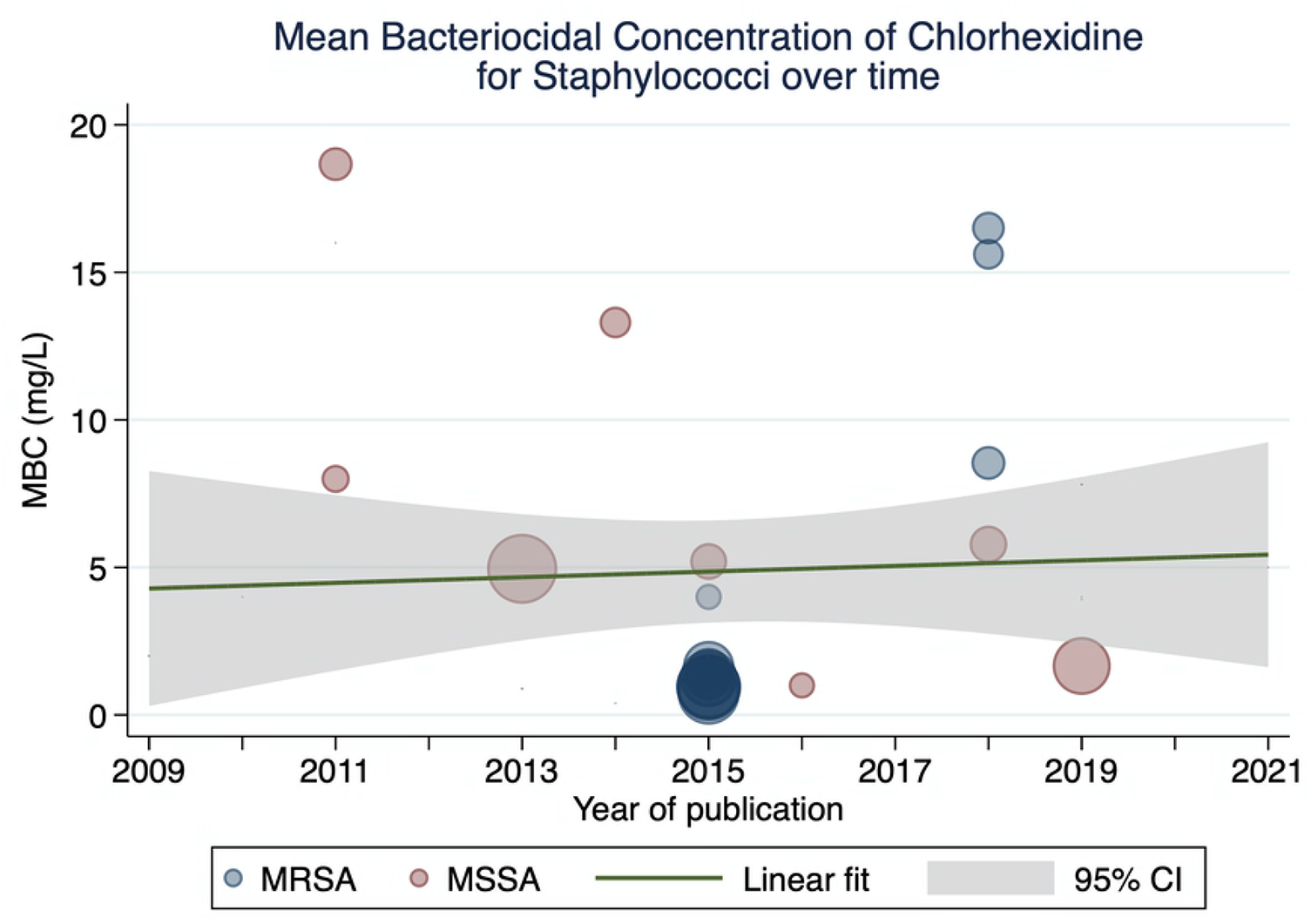
A scatterplot of study-level estimates of mean MBC over time, for *Staphylococci*. The size of the points corresponds to the precision (inverse variance) of the study.

Overall, 25 studies reported the MBC of CHX for *Streptococci* species; observations of viridans *Streptococci* were most common(44,45,47,48,52–54,58,66,69,75,77–79,81,82,84–86,89,99,100,103,105,106) and 1 study(106) provided an estimate for *Streptococcus pyogenes* (Lancefield group A). The pooled mean MBC of CHX for *Streptococci* was 8.54 mg/L (95% CI 4.75, 12.3; I^2^ 99%). Meta-regression showed that the MBC of CHX for *Streptococci* had not changed over time (β 0.13 [-0.35, 0.62]; I^2^ 97%; Fig 4).

**Fig 4.**
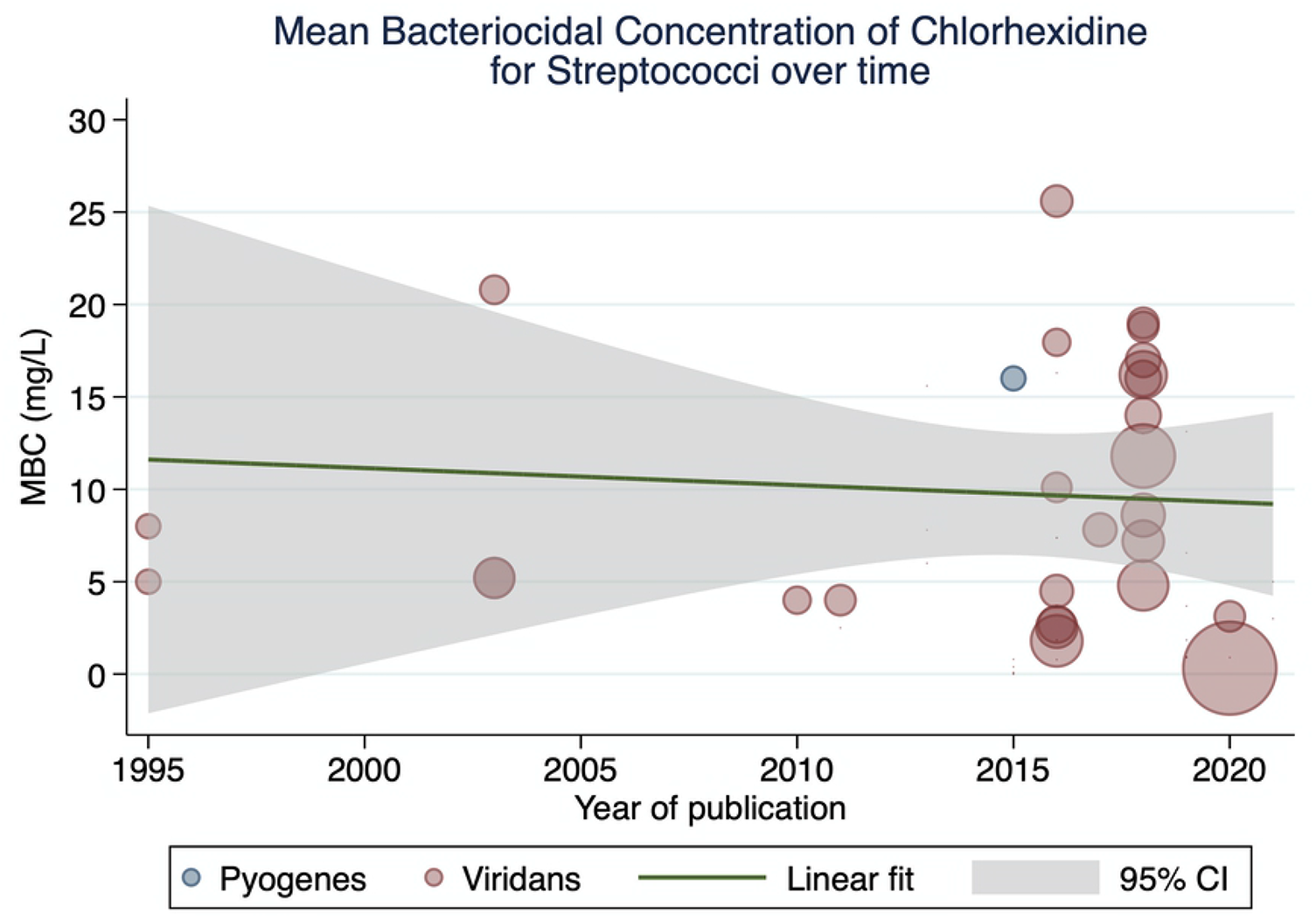
A scatterplot of study-level estimates of mean MBC over time, for *Streptococci*. The size of the points corresponds to the precision (inverse variance) of the study.

There were insufficient data for meta-analysis of the MBC of PVI. Also, the majority of MBC data for PVI was derived from studies of Enterobacteriales, which is not a common cause of surgical site infection and so equally, not clinically relevant.

## Discussion

This review summarise the evidence to-date and suggests that there has been no increase in MBC of CHX for the main SSI-causing skin microbes in recent decades. The stability of CHX susceptibility is reassuring for clinicians and policy makers alike, as it endorses current surgical guidance worldwide advocates topical alcoholic chlorhexidine for skin asepsis prior to surgery.

The definition of bacterial susceptibility and resistance to biocides is still a matter of debate. Most clinically relevant bacteria have defined susceptibility and resistance to systemic antibacterials based upon the MIC relative to an epidemiologically derived clinical breakpoint. However, the use of MICs is less useful in determining the efficacy of topical antiseptics biocides given the desire to induce death of specific microbes rather than inhibition. Understanding the lethality of a biocide (hence MBC) is therefore a potentially more attractive measure than MIC for topical antiseptics(114). Additionally, the relevance of both MICs and MBCs with respect to biocides has been questioned.(15) Chlorhexidine for skin prep is used in concentrations of 5000 to 50000 μg/mL and with MBC values ranging from 0 to 30 μg/mL, this is one thousand times greater than the apparent required concentration(115).

MIC and MBC rely on attaining a steady concentration in bodily fluids as seen by the pharmacodynamics of antibiotics(116). The EUCAST definition of a susceptible organism is “a microorganism is defined as susceptible by a level of antimicrobial activity associated with a high likelihood of therapeutic success”(117). Therapeutic success in the context of topical CHX prior to surgery is disinfection; complete elimination of all relevant micro-organisms, except certain bacterial spores(118). Measurement of MIC and MBC are gained from *in vitro* susceptibility testing of microorganisms to topical biocides and therefore, provide little information as to the mechanism of resistance or likely clinical outcome. Therefore, our recommendation is the utilisation of epidemiological cut-off values (ECOFF) based on MBC distributions to better understand the response of bacteria to CHX in clinical practice. An analysis of ECOFF values of CHX to common bacterium, including those commonly causing SSI did not reveal a bimodal distribution, concluding that resistance is uncommon to CHX in natural populations of clinically relevant organisms(88).

While values above a certain cut off may be defined as a breakpoint and hence resistant, this needs to be correlated with the clinical picture. Does CHX still achieve adequate disinfection in a population of bacterium with a MBC value greater than the 95% ECOFF? Without this information, our understanding of how MBC values beyond the normal distribution impacts clinical use remains poor. However this is contrary to the application of biocides in clinical settings prior to surgery, particularly during skin preparation of sensitive areas (e.g. the face) and in the presence of open wounds(6). We speculate that real-time methods of quantifying bacterial colonisation (e.g. free bacterial DNA/RNA sequency tools such as MiniION), taken before and serially after topical application of CHX, may be more clinically useful in understanding the true bactericidal effect of CHX, what microbes is acts upon preferentially, how long it exerts its action and how this is achieved.

## Limitations

Most of the included studies reported MIC rather than MBC, which meant that much data could not be synthesised. This might explain why our meta-data (for *Staphylococci* and *Streptococci* at least) disagrees with individual articles(18,19,21,22). Papers also often failed to report the type and concentration of chlorhexidine used which hindered meta-regression. Some papers clearly stated the year that the microbial strain was isolated, although this was often unclear and therefore the date of the paper was taken as the year of the isolate which may not adequately represent the change in MBC over time. This was also the case with location, so where possible the location of the laboratory or hospital was used as a surrogate. Isolates from a clinical setting are exposed to different selection pressures. As 61% of the microbes used were laboratory isolates as opposed to clinical isolates, it is unclear how our data can be generalised to clinical environments.

## Conclusion

There has been no demonstrable change in the susceptibility of surgical site infection causing pathogens to chlorhexidine over time. A clear definition of reduced susceptibility and resistance of pathogens to biocides is needed, alongside consensus on the methods for measuring these phenomena.

## Contributions

RA and VHD contributed equally to this paper and are joint first authors. They co-designed the protocol, extracted data and co-authored the manuscript.

CJ provided clinical oversight for this study, supporting the design, advising on data extraction, analysis techniques and co-authored the manuscript.

RGW conceived the idea of this study, co-designed the protocol, supervised all elements of the delivery, extracted, and checked data, performed the statistical analysis and co-authored the manuscript

## Notes

### Competing Interest Statement

The authors have declared no competing interest.

